# Long-term effects of early postnatal stress on Sertoli cells functions

**DOI:** 10.1101/2022.07.27.501498

**Authors:** Kristina M. Thumfart, Samuel Lazzeri, Francesca Manuella, Isabelle M. Mansuy

## Abstract

Sertoli cells are somatic cells in testes essential for spermatogenesis, as they support the development, maturation, and differentiation of germ cells. Sertoli cells are metabolically highly active and physiologically regulated by external signals, particularly factors in the blood stream. In disease conditions, circulating pathological signals may affect Sertoli cells and consequentially, alter germ cells and fertility. While the effects of stress on reproductive cells have been well studied, how Sertoli cells respond to stress remains poorly characterized. Therefore, we used a mouse model of early postnatal stress to assess the effects of stress on Sertoli cells. We developed an improved enrichment strategy based on intracellular stainings and obtained enriched preparations of adult Sertoli cells from exposed males. We show that adult Sertoli cells have impaired electron transport chain (ETC) pathways and that several components of ETC complexes I, III, and IV are persistently affected. We identify the circulation as a potential mediator of the effects of stress, since treatment of primary Sertoli cells with serum from stressed males induces similar ETC alterations. These results newly highlight Sertoli cells as cellular targets of early life stress, and suggest that they may contribute to the negative effects of stress on fertility.

**Highlights:**

- We present an improved method to obtain enriched preparations of Sertoli cells from adult mouse testis for molecular analyses
- Sertoli cells from adult males exposed to stress during early postnatal life have altered electron transport chain (ETC) expression, suggesting persistent effects of early life stress on Sertoli cells physiology
- Serum from adult males exposed to early postnatal stress reproduces ETC gene dysregulation in cultured Sertoli cells.

**Graphical abstract:** 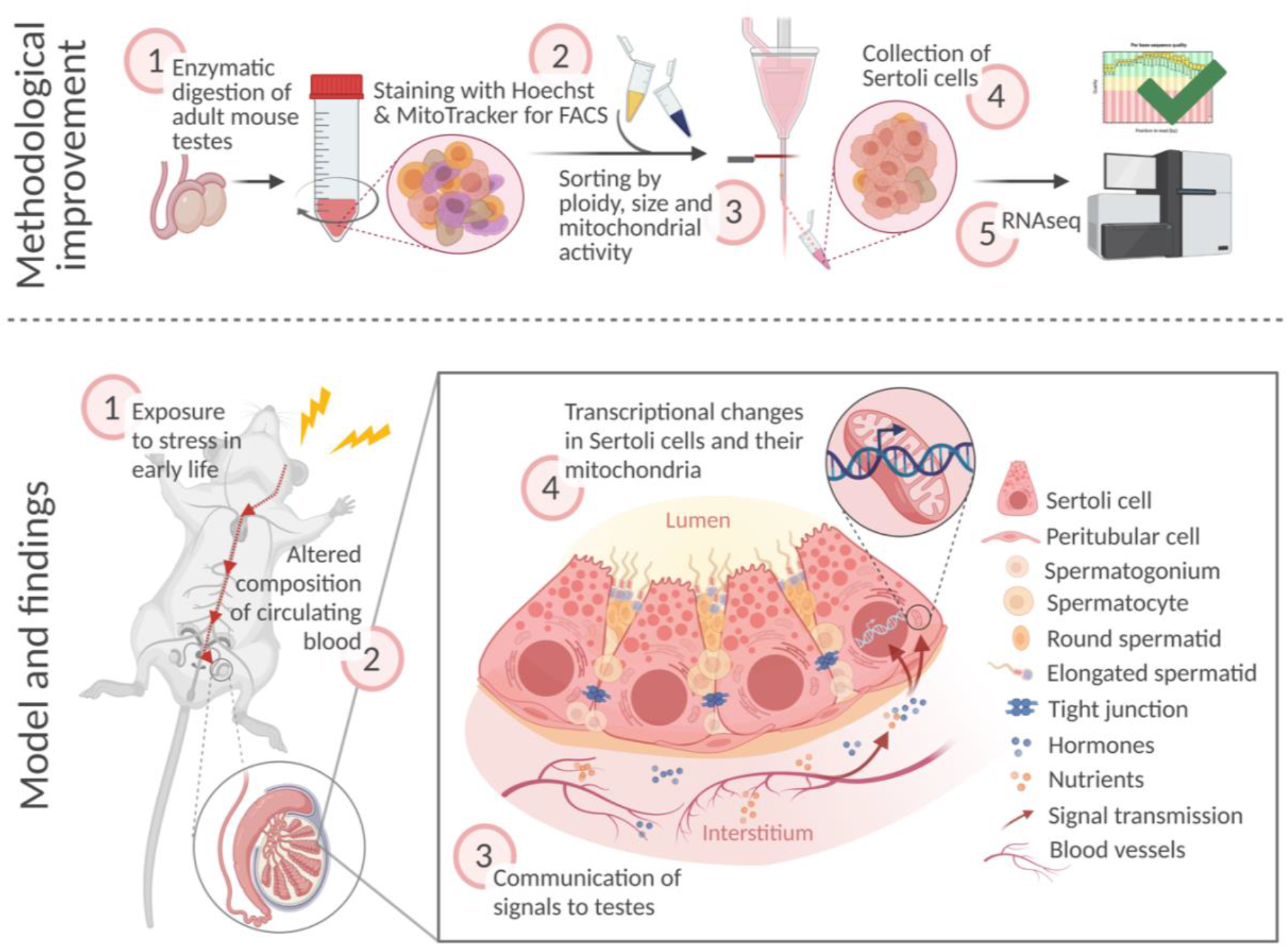

## Introduction

Sertoli cells are somatic cells in the seminiferous tubules of testes tightly associated with germ cells and essential for spermatogenesis. They provide physical and structural support to differentiating spermatogenic cells and form and maintain a protective blood-testis barrier (Griswold 2018). Sertoli cells have paracrine functions and secrete growth factors, hormones, cytokines, and extracellular vesicles (Mancuso et al. 2018). These factors provide developmental guidance and immunological protection to germ cells (Mäkelä and Hobbs 2019; Kaur et al. 2020). Sertoli cells have a high glycolytic flux to provide nutritional support for germ cells. Through glycolysis, they metabolize glucose into lactate, which is the primary source of energy for spermatocytes and spermatids (Zhang et al. 2018). For their own energy needs, Sertoli cells rely on oxidative phosphorylation of lipids, which they receive through the blood stream or through the recycling of germ cell waste material (Regueira et al. 2018). Oxidative phosphorylation is catalyzed by four complexes of the electron transport chain (ETC) located in the mitochondrial inner membrane. These complexes use energy generated from nutrient oxidation to create a proton gradient across the mitochondrial inner membrane, which is then used by the ATP-synthase (complex V) to generate ATP (Nolfi-Donegan, Braganza, and Shiva 2020).

Sertoli cells are in close contact with blood vessels to sense hormones and metabolites present in the blood stream, and thereby receive signaling from circulating factors (Rebourcet et al. 2016). Changes in circulating factors in pathological conditions may therefore alter Sertoli cell metabolism and physiology and affect spermatogenic cells. This is particularly critical in early life, because Sertoli cells lose their mitotic activity during postnatal development and thus, if they are affected in early life, they are likely to remain so until adulthood (Sharpe et al. 2003). Indeed, altered blood homeostasis due to neonatal hormonal dysregulation in mice (Sarkar and Singh 2017) or early exposure to environmental toxins in rats (de Oliveira et al. 2020; Sadler-Riggleman et al. 2019) were shown to alter the energy metabolism of Sertoli cells. Exposure to high fat diet and resulting diabetes can also alter both glucose and lipid metabolism of mouse Sertoli cells (Luo et al. 2020), which may contribute to altered reproductive functions in response to diabetes (Sajadi et al. 2019).

To gain insight into the effects of early life stress on Sertoli cells, we examined the transcriptome of Sertoli cells from adult males exposed to stress in early postnatal life using an improved method to enrich Sertoli cells from adult mouse testes. We observed that oxidative phosphorylation by the mitochondrial ETC is altered in adult Sertoli cells, and that many ETC components are affected. We further show that serum can recapitulate ETC components alterations in cultured Sertoli cells, suggesting the involvement of circulating factors in the alterations.

## Results

### Enrichment of Sertoli cells from adult testis

To obtain Sertoli cells from adult mouse testis, we developed an enrichment method based on fluorescence-activated cell sorting (FACS) not requiring any transgenic or surface marker (Fig 1a). First, testis tissue is digested sequentially in collagenase, trypsin and hyaluronidase (Bhushan et al. 2016), then cells are processed through FACS. While collagenase digests the interstitium and detaches seminiferous tubules from each other, trypsin fragments tubules and detaches peritubular cells. Hyaluronidase separates Sertoli cells from germ cells. The FACS strategy is based on intracellular staining with Hoechst and MitoTracker based on specific properties of Sertoli cells including diploidy (post-mitotic state) (Sharpe et al. 2003), large size (Wong and Khan 2021), and high metabolic activity compared to other cells in testes (Miettinen and Björklund 2017). Plotting of Hoechst intensity versus forward scatter (FSC), indicating size, identified several testicular subpopulations (Fig 1b). Diploid cells were separated from haploid (spermatids) and tetraploid (dividing) cells distinguished by Hoechst intensity (Fig 1c) (Gaysinskaya et al. 2014). Diploid cells were then fractionated by size using high and low FSC (Fig 1d), and cells of the high FSC fraction were separated into high and low MitoTracker signal (high/low APC) for mitochondrial mass and activity (Fig 1e) (Clutton et al. 2019). Vimentin staining of single-cell suspension collected from seminiferous tubules before FACS identified 1.9 ± 0.6% (weighted mean ± weighted standard deviation) of Sertoli cells (Fig 1f) and after FACS, 14.8 ± 3.6% in the fraction of diploid cells (Fig 1g). This was further increased to 37.7 ± 30% in the high FSC cells fraction (Fig 1h) and up to 89.8 ± 5% in the fraction of cells with a high APC signal (Fig 1i).

**Fig 1.**
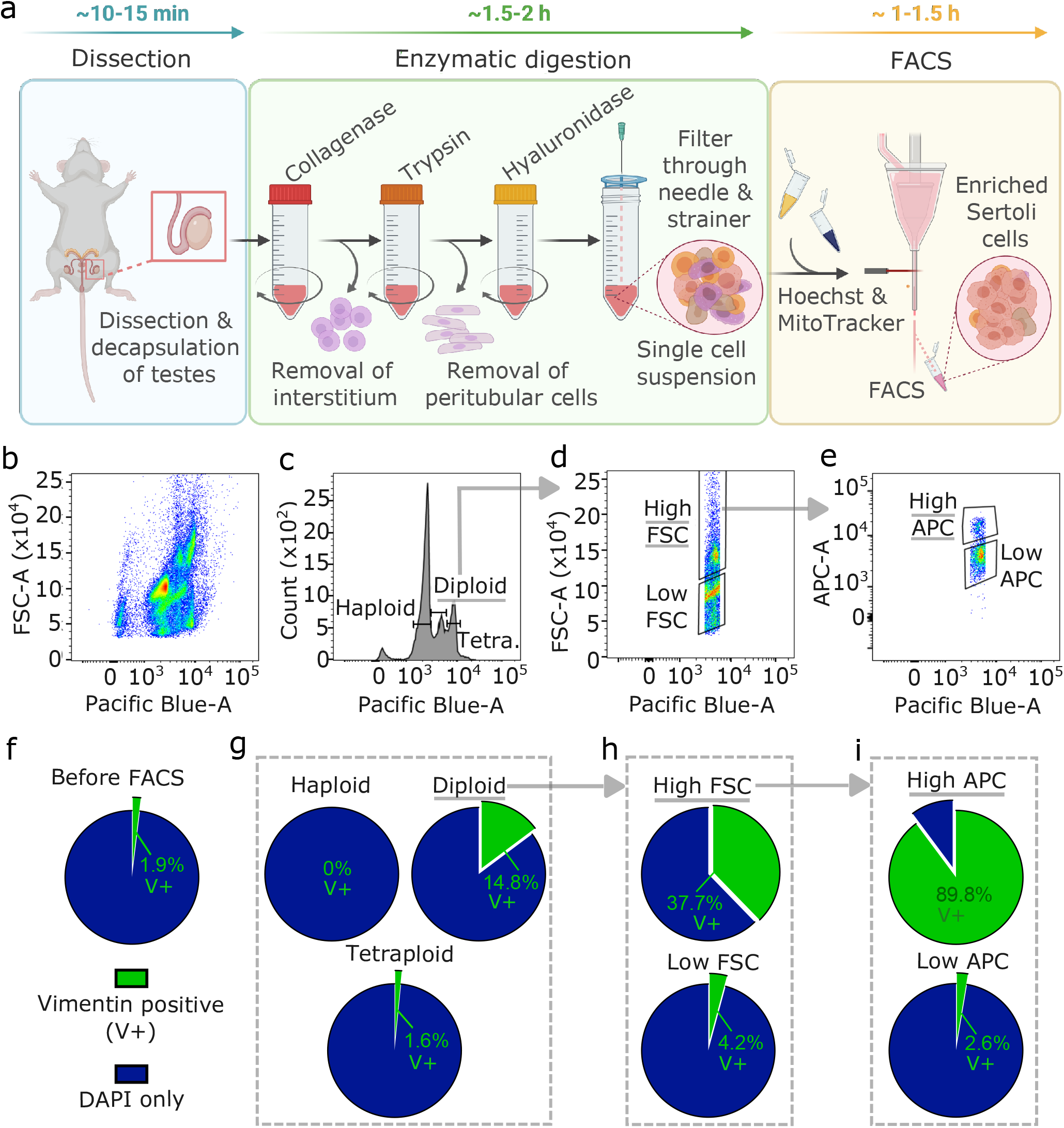
Sertoli cells enrichment and visualization by vimentin staining. (a) Workflow for Sertoli cells enrichment procedure including testis dissection, enzymatic digestion by collagenase, trypsin, and hyaluronidase, followed by staining with Hoechst and MitoTracker for FACS. Time estimates are indicated for the individual steps. (b-e) Results of FACS profiles during Sertoli cells enrichment. (b) Hoechst signal (Pacific Blue) plotted against FSC (indicating size) showing several cell populations in the single cell testis preparation. (c) Diploid cells selected using the Hoechst signal, (d) large cells using a high FSC signal, then (e) cells with high mitochondrial activity using MitoTracker (high APC) signal. (f-i) Enrichment of Sertoli cells (vimentin-positive, V+) in different FACS fractions. Percentage of V+ cells of all cells (DAPI) is shown (f) before FACS, (g) after selection of diploid cells, (h) after size selection by high FSC within diploid cells, and (i) after gating on high mitochondrial activity within diploid, high FSC cells (weighted averages of 4 independent replicates). Counts are summarized in S1 Table. Underlined fractions in (b-i) correspond to gates chosen for Sertoli cells enrichment. Arrows indicate implementation of gate settings before further partitioning into sub-gates.

### Transcriptomic profiling of enriched Sertoli cells by RNA sequencing

We characterized the transcriptome of the Sertoli cells enriched from adult mouse testis by RNA sequencing and examined known markers of testicular cell populations using published single-cell sequencing datasets (Green et al. 2018). We observed that several Sertoli cell markers including *Amhr2, Clu, Ctsl, Rhox5*, and *Sox9* were more abundant in the isolated cells than markers of other testicular cells, validating the enrichment protocol (Fig 2a). The top 100 expressed genes were screened for enriched *Kyoto Encyclopedia of Genes and Genomes* (KEGG; Fig 2b) and *Gene Ontology* (GO; Fig 2c) pathways. Identified pathways involve immunological regulation, energy metabolism, cell-cell junctions, phagocytosis, and secretion, consistent with known Sertoli cell functions (Griswold 2018).

**Fig 2.**
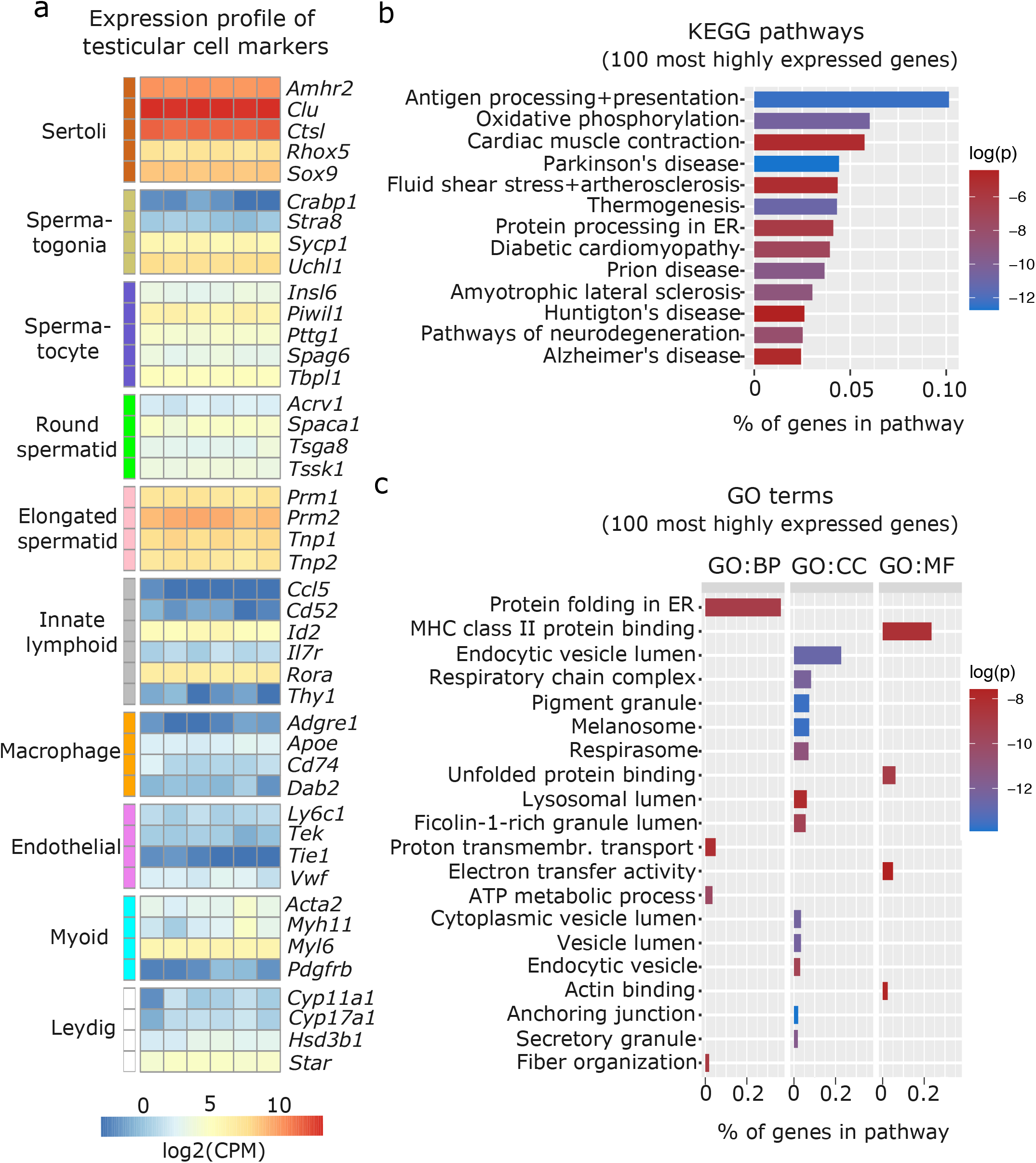
Characterization of enriched Sertoli cells by RNA sequencing. (a) Heatmap of testicular cell markers in enriched Sertoli cells, ordered by cell type-specificity including Sertoli cells, spermatogenic cells (spermatogonia, spermatocytes, round and elongated spermatids), immune cells (lymphoid and macrophage), endothelial cells, smooth muscle cells (peritubular myoid) and endocrine cells (Leydig). Color scale indicates normalized log2 gene counts per million (CPM). Enriched (b) KEGG and (c) GO pathways for 100 most highly expressed genes in collected Sertoli cells including immunological (e.g. KEGG: Antigen processing and presentation; GO MF: MHC class II protein binding), metabolic (e.g. KEGG: Oxidative phosphorylation, GO CC: respiratory chain complex), cell-cell junction (KEGG: Cardiac muscle contraction, GO, CC: anchoring junction), phagocytosis (GO, CC: endocytic vesicle lumen, lysosomal lumen), and secretion (KEGG: protein processing in ER, GO, CC: secretory granule) pathways. Ratio of genes per pathway is given on the x-axis and log of p-value (log(p)) is indicated on a color scale. BP: biological process, CC: cellular component, MF: molecular function.

### Persistent changes in Sertoli cells transcriptome caused by early life stress

We used our improved enrichment method to obtain Sertoli cells from adult males exposed to stress in early postnatal life and examined the effects on the cells. As a stress paradigm, we used an established mouse model of postnatal stress based on unpredictable maternal separation combined with unpredictable maternal stress (MSUS) (Franklin et al. 2010). Newborn pups were separated from their mother unpredictably 3 hours each day from postnatal day 1 to 14 (PND1-14) and during separation, mothers were stressed unpredictably. This paradigm induces persistent metabolic and behavioral alterations in exposed animals when adult, and in their progeny across several generations (Gapp et al. 2014; Franklin et al. 2010). We collected Sertoli cells from adult MSUS and control mice from two independent cohorts (batch 1 and 2) and profiled their transcriptome by RNA sequencing in batch 1, followed by validation by quantitative PCR in batch 2 (Fig 3a). Using over-representation analyses of all genes with a p-value <0.05 from the RNA sequencing datasets, we observed that the most significantly altered molecular pathways (top five) were related to the mitochondrial ETC (S2 Table). Among KEGG pathways, “oxidative phosphorylation” (p<0.001) was the most significant (Fig 3b), while among GO terms for cellular components, “respirasome” (p<0.001) and “respiratory chain” (p<0.001) were the most significantly enriched (S1a Fig). To visualize ETC gene expression compared to the general expression distribution of all genes in the RNA sequencing datasets, we plotted candidates of the “oxidative phosphorylation” pathway with a p-value of <0.05 for control Sertoli cells (Fig 3c). All ETC components are more highly expressed than the average gene in Sertoli cells. Genes encoded by mitochondrial DNA (*ND2, ND4, ND6, CYTB, COX1, COX2*) have higher, but more variable expression than genes encoded by nuclear DNA (Fig 3c). This is consistent with recent single-cell RNA sequencing datasets in mouse testis (Green et al. 2018). In Sertoli cells from MSUS males, ETC genes encoded by nuclear DNA were primarily downregulated compared to controls (Fig 3d), while changes in mitochondrial genes were more variable (S2a Fig). Validation of changes in ETC genes in a second batch of Sertoli cells by multiplex RT-qPCR screen (Fluidigm) confirmed that the majority of ETC genes were downregulated in MSUS Sertoli cells (Fig 3e). Out of 18 target genes identified from the KEGG pathway “oxidative phosphorylation”, 4 were downregulated significantly (p<0.05; *ND4, ND6, CYTB, Uqcr11*), while no genes were significantly upregulated. When looking closer at downregulated genes, particularly components of ETC complex I, III, and IV were downregulated, while components of complex II and V had variable expression changes. Also, mitochondrial genes were predominantly downregulated, which was different to what we expected from the RNA sequencing data of batch 1, which showed higher inconsistency in mitochondrial gene expression changes (S2a Fig). However, this bias might be attributable to significant mitochondrial copy number variation (CNV) in MSUS vs. control Sertoli cell samples of batch 1 (S2b Fig).

**Fig 3.**
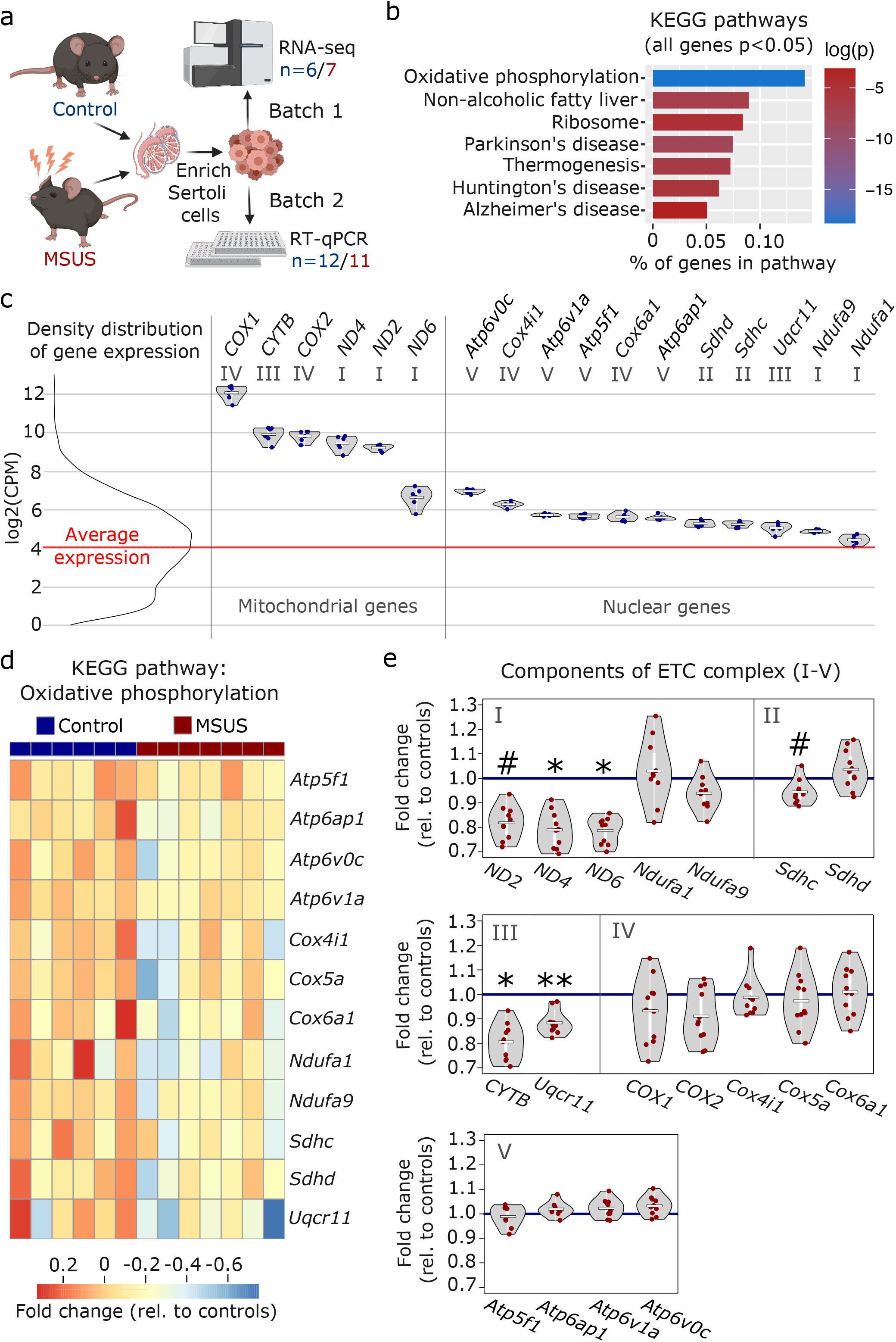
Transcriptomic analyses of MSUS and control adult Sertoli cells. (a) Schematic representation of the experimental strategy for Sertoli cells enrichment and transcriptomic analyses for control (blue) and MSUS (red) mice. For RNA sequencing, n=6 controls and n=7 MSUS (Batch 1). For validation of candidate genes with RT-qPCR, n=12 controls and n=11 MSUS (Batch 2). (b) KEGG pathways of significantly altered genes (p<0.05) in Sertoli cells from MSUS males with % genes per pathway (x axis) and log of p-value (log(p)) on vertical color scale. (c) Expression profile of ETC candidate genes from KEGG pathway “oxidative phosphorylation” in control samples (scale is in log2(CPM)). Left, density distribution of all expressed genes, with red line indicating the average expression of genes in control Sertoli cells. Right, candidate genes encoded by mitochondrial (Mitochondrial genes) and nuclear (Nuclear genes) DNA plotted on the same scale. Roman numbers indicate name of ETC complex. Mean of control samples depicted as white line, individual samples as blue dots. (d) Heatmap of nuclear encoded genes with p<0.05 of KEGG pathway “oxidative phosphorylation”. Fold change relative to controls is indicated in the color scale. (e) RT-qPCR of ETC candidate genes in batch 2 samples. Candidate genes are divided according to ETC complexes (I, II, III, IV, V) and fold change of expression profiles is shown for MSUS samples relative to control mean (blue line) of respective genes. Mean of MSUS samples depicted as white line, individual samples as red dots. **p<0.01, *p<0.05, #p<0.1, student’s t-test.

### Serum from MSUS males downregulates ETC components in primary Sertoli cells

Our previous work showed that serum from MSUS males can induce molecular changes in reproductive cells, when injected intravenously to adult males or when used to treat immortalized spermatogonial stem cells (van Steenwyk et al. 2020). Since Sertoli cells receive signals from the blood stream, we examined if serum from MSUS males can reproduce changes in ETC components observed after MSUS. We prepared primary Sertoli cell cultures from mouse testis and supplemented them with 10% serum obtained from MSUS or control mice for 24 hours (Fig 4a). Analyses of candidate genes by RT-qPCR showed that 4 ETC components are significantly downregulated (p<0.05; *ND6, Ndufa1, COX1, Cox6a*) (Fig 4b). Downregulation was most consistent for components of complex I, III, and IV while complex II and V components were more variably altered, and some of them upregulated (p<0.05; *Sdhc, Atp6ap1*).

**Fig 4.**
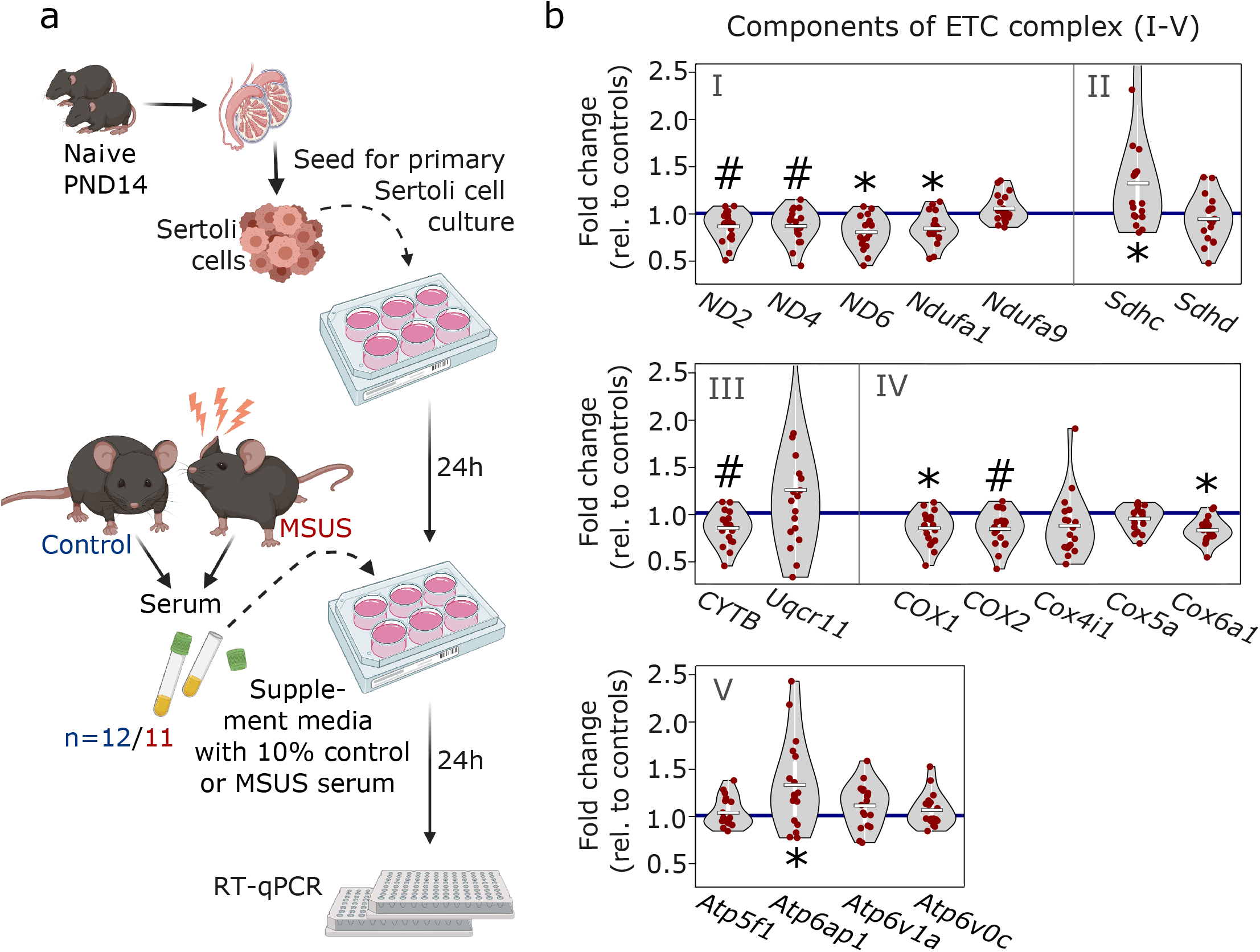
Analyses of ETC genes in primary Sertoli cells after MSUS serum exposure. (a) Schematic representation of the experimental strategy for serum treatment of Sertoli cells. Primary Sertoli cells from naïve mouse pups (PND14) were seeded and treated with 10% serum from control (blue) and MSUS (red) mice for 24h. Treated cells were then harvested for RT-qPCR. Controls, n=12; MSUS, n=11. Experiment was performed in duplicates. (b) RT-qPCR of ETC candidate genes in serum treated Sertoli cells. Candidate genes are divided according to ETC complexes (I, II, III, IV, V) and fold change of expression profiles is shown for MSUS samples relative to control mean (blue line) of respective genes. Mean of MSUS samples depicted as white line, individual samples as red dots. *p<0.05, #p<0.1, Kenward Roger method

### Exposure to MSUS serum does not alter metabolic functions of primary Sertoli cells

Differential regulation of the ETC can affect the intracellular redox state of cells and alter their lactate-pyruvate ratio (Titov et al. 2016; Patgiri et al. 2020). Therefore, we examined if the downregulation of ETC complex I, III, and IV components has metabolic consequences for Sertoli cells. We measured lactate and pyruvate in the conditioned medium of primary Sertoli cells after MSUS serum treatment and examined ROS activity by 2’,7’–dichlorofluorescin diacetate (DCFDA) staining. The level of lactate and pyruvate was not significantly altered (S3a,b Fig). Likewise, the ratio of lactate to pyruvate (S3c Fig) or ROS levels (S3d Fig) were not altered. These results suggest that the downregulation of ETC complexes I, III, and IV after MSUS serum exposure was not sufficient to affect metabolic functions of Sertoli cells.

## Discussion

This study examines the effects of early life stress on somatic cells in the adult mouse testis and addresses the question of which factors may play a role in the induction of these effects. Using an established mouse model of stress, we show that Sertoli cells from adult males exposed to stress in early postnatal life have altered ETC pathways. The alterations affect several mitochondrial complex components, which are predominantly downregulated in adult Sertoli cells. We link these alterations to circulating blood factors by showing that the ETC complexes downregulation can be reproduced in primary Sertoli cells in culture when the cells are treated with serum from exposed adult males. These results suggest that Sertoli cells can be persistently altered by adverse conditions in early life and keep a biological trace of exposure for many months. This may be explained by the fact that these cells are post-mitotic in the adult testis and are no longer able to self-renew unlike spermatogenic cells. They therefore do not have the possibility to correct or erase molecular changes by cell division and renewal, and remain altered until adulthood, possibly throughout life. Other environmental factors such as endocrine disruptors have also been found to affect Sertoli cells in rats (Guerrero-Bosagna et al. 2013).

Since Sertoli cells are essential for germ cell maintenance and physiology, their persistent alterations during development through to adulthood may affect spermatogenesis and have detrimental consequences for germ cells and fertility. Psychological stress has indeed been reported to reduce fertility in humans (Bräuner et al. 2020) and is known to lead to molecular changes in spermatogenic cells in testis (Tian et al. 2021) and adult sperm in rodents (Gapp et al. 2014; Franklin et al. 2010), with the potential to impair metabolism and behavior in the offspring (Gapp et al. 2014). However, the mechanisms by which Sertoli cells may alter germ cells are not known.

Our data that ETC components in Sertoli cells are affected, suggest a link between stress exposure at a young age and mitochondrial functions in the adult. Mitochondria are organelles known for their ability to adjust to changes in metabolic demand in cells (Bereiter-Hahn and Vöth 1994). Thus, they are sensitive targets of systemic cellular perturbations and potential sensors of environmental exposure. Indeed, ETC components in brain and muscle have already been shown to be altered by early postnatal stress in mice (Ruigrok et al. 2021). Our data extend these findings by showing that ETC complexes are also affected persistently in Sertoli cells by early postnatal stress, and provide candidate molecular targets to examine in relation to potential germ cell damage. The downregulated complexes I, III, and IV have in common to be able to transport protons across the inner mitochondrial membrane, and contribute to the generation of a proton gradient for ATP production (Marreiros et al. 2016). This could influence the metabolism of not only Sertoli cells, but also of neighboring germ cells via altered extracellular signaling pathways such as redox state and lactate production (Titov et al. 2016; Patgiri et al. 2020). However, in cultured Sertoli cells, reproducing ETC pathways alterations with serum from stressed males did not affect reactive oxygen species (ROS), lactate or pyruvate level. This suggests that changes may only be subtle and cannot be detected by classical methods such as fluorescent assays *in vitro*, or may be compensated for by alternative mechanisms. Using sensitive substrate sensors such as genetically encoded fluorescence resonance energy transfer (FRET) sensors to detect metabolite flows *in vitro* or *in vivo* may help identify changes (Mächler et al. 2016). Other systemic effects by cell-cell-communication within testis or signaling through innervation and via the lymphatic system may also occur.

Classically, methods to enrich Sertoli cells are based on specific culture procedures and conditions which have some limitations. For instance, *Datura stramonium* (DSA)-lectin coated dishes can be used to favor the attachment of Sertoli cells (Scarpino et al. 1998) and allow easier removal of contaminating germ cells by washing and/or hypotonic shock (Wagle et al. 1986; Anway et al. 2003). However, culture conditions can introduce biases to cells and modify their epigenetic landscape and functions compared to *in vivo* (Zomer and Reddi 2020a). Therefore, enrichment methods not requiring any culture, but allowing to isolate cells directly from tissue, are advantageous for molecular analyses. For Sertoli cells, transgenic or knock-in mice expressing fluorescent proteins under the control of *Amh* or *Sox9* promoters have been generated and can yield relatively pure Sertoli cell preparations by FACS (Zimmermann et al. 2015; Zomer and Reddi 2020b). However, wildtype mice may be preferable to avoid possible transgene interference (in homozygous mice for instance) or GFP protein toxicity, and for easier availability without requiring any specific breeding scheme. This is particularly needed for large-scale *in vivo* experiments that require big cohorts for phenotyping like behavioral, physiological and/or metabolic testing. Our FACS-based method provides an efficient alternative through capitalizing on previous work in fixed cells (Rotgers et al. 2015), using parameters that separate testicular populations by ploidy through DNA staining and light scattering via cytometry. Due to intracellular stainings with Hoechst and MitoTracker, biases due to cleavage or internalization of surface antigens after enzymatic digestion can be avoided (Autengruber et al. 2012; Tsuji et al. 2017). Using this method, we obtain a high enrichment of Sertoli cells confirmed by vimentin staining (Fig 1 f-i) and specific markers expression in obtained cells (Fig 2a). Notably, markers of elongated spermatids such as *Prm2* were detected in our Sertoli cells datasets, similarly to previously reported in testis single-cell sequencing datasets (Green et al. 2018). These marker transcripts likely correspond to remnants of spermatids phagocytosed by Sertoli cells that persist in their cytoplasm. Lastly, we cannot exclude that Hoechst and MitoTracker binding affects DNA and mitochondria integrity in sorted Sertoli cells. However, incubation with the stains is kept to a minimum and cells are placed on ice at all times after staining.

In conclusion, our findings highlight the vulnerability of Sertoli cells during postnatal development and the fact that they can be persistently altered by stress exposure. Whether and how this may ultimately affect germ cells functions and physiology is still an open question that needs to be investigated.

## Methods

### Animals

Adult C57Bl/6J mice (3-5 months old) were kept under a 12-h reverse light/dark cycle in a temperature- and humidity-controlled facility with access to food and water ad libitum. All experiments were performed during the active (dark) cycle of the mice in accordance with guidelines and regulations of the Cantonal Veterinary Office, Zürich (animal licenses ZH057/15 and ZH083/18).

### MSUS paradigm

3-month old female and male breeders were randomly paired and assigned to MSUS or control groups. Newborn pups in the MSUS group were separated from their mother for 3 h per day at unpredictable times from postnatal day (PND) 1 to 14. Any time during separation, mothers underwent an unpredictable acute swim in cold water (18°C for 5 min) or 20-min restraint in a tube. Control animals were left undisturbed. Pups were weaned at PND21 and assigned to new cages according to group and gender (3–5 mice/cage). Siblings were assigned to different cages to avoid litter effects. An overview of MSUS and control mice used for tissue collection is presented in S3 Table.

### Testis collection

Adult mice were single-housed with food and water *ad libitum* the night before sacrifice to minimize stress. Mice were sedated with isoflurane before decapitation. For testis collection in pups at PND14, whole litters (4-6 males on average) were sacrificed by decapitation soon after being removed from their mother.

### Enzymatic digestion of mouse testis

Testes were dissected, decapsulated and placed into a 50 ml canonic tube containing 10 ml of enriched DMEM/F12 medium (1x DMEM/F12 [Gibco], supplemented with 15mM HEPES, 1x GlutaMAX [Gibco], 1x Minimum Essential Medium Non-essential Amino Acids [Gibco] and 1% penicillin-streptomycin [Pen-Strep; Gibco, 10,000 U/ml]). The tissue was transferred to 5 ml collagenase solution (1mg/ml collagenase [from Clostridium histolyticum, Sigma Aldrich] and 0.02 mg/ml DNase [from bovine pancreas, Sigma Aldrich] in enriched DMEM/F12) and incubated at 35°C for 5-10 minutes with intermittent shaking until seminiferous tubules dissociated from the interstitium. For washing, 25 ml of enriched DMEM/F12 were added, the tube was inverted three times and tubules were allowed to settle for 2-3 minutes. Supernatant containing interstitial cells was discarded and the washing step was repeated twice. Then, 5 ml 0.25% trypsin-EDTA solution (Gibco) supplemented with 0.1 mg/ml DNase were added to the tubules and incubated at 35°C for 5-10 minutes with intermittent shaking until tubules were fragmented. Tubules were washed one time with enriched DMEM/F12 containing 10% fetal bovine serum (FBS; HyClone, Cat. No. SV30160.03) to inactivate trypsin and were allowed to settle for 5 minutes. Supernatant containing peritubular cells was removed and washing was repeated with enriched DMEM/F12 two more times. To obtain a single-cell suspension from the cleaned seminiferous tubules, the tissue was further digested in hyaluronidase solution (1mg/ml hyaluronidase [from sheep testes, Sigma Aldrich] and 0.02 mg/ml DNase in enriched DMEM/F12) for 5-10 more minutes at 35°C with intermittent shaking. For proper dissociation of cells, they were passed through a 5 ml serological pipette 4-5 times, then 25 ml enriched DMEM/F12 were added. Cells were centrifuged at 400xg for 3 minutes, the supernatant was removed and cells were resuspended in 10 ml enriched DMEM/F12. To remove any remaining cell clumps, the cell suspension was slowly passed through a 20G needle, then filtered through a 70 µm cell strainer. 25 ml of enriched DMEM/F12 were added and cells were centrifuged at 400xg for 3 minutes and collected for further enrichment.

### Blood processing

To obtain serum, trunk blood was collected and allowed to clot for 15-30 minutes at room temperature (RT). To separate serum from the clot, samples were centrifuged for 10 minutes at 2,000 x g. The supernatant (serum) was transferred to a new tube and stored at -80°C until further use.

### Fluorescence-activated cell sorting (FACS)

Cells obtained after enzymatic digestion of testis were resuspended in 5-10 ml FACS buffer (1x DPBS [Gibco] supplemented with 1% Pen-Strep, 1% FBS, 10 mm HEPES, 1 mm pyruvate [Gibco] and 1 mg/ml glucose [Gibco]) and counted with a hemocytometer. Cells were diluted at 10^6^/100 µl in FACS buffer. 1 µl of Hoechst 33342 Solution (BDPharmingen, stock: 1 mg/ml) and 0.1 µl of MitoTracker Deep Red (Invitrogen, stock: 1mM) were added per 100 µl cell suspension, then cells were incubated at 35°C for 20 minutes. Thereafter, cells were kept on ice at all times. Cells were washed twice with ice-cold FACS buffer and the Sertoli cell fraction was sorted according to the FACS diagram depicted in Fig 1b-e. Briefly, cell debris and doublets were gated out and remaining cells were gated for diploidy using the Hoechst channel. Diploid cells were further gated for high FSC and subsequently for a high signal in the MitoTracker channel.

### Immunocytochemistry

Round coverslips (diameter: 8mm, thickness: 1, Warner Instruments) were placed into 48-well plates and coated with Poly-L-lysin solution (P8920, Sigma-Aldrich) for at least 15 min at RT. Coverslips were then washed three times with distilled, autoclaved water and were allowed to dry overnight. The day after, always 40,000 cells in enriched DMEM/F12 medium supplemented with 10% FBS were plated onto the slides and allowed to attach at RT for at least 20 min and another hour in an incubator at 37°C. Thereafter, medium was aspirated and cells were fixed with 4% paraformaldehyde (PFA) for 15 min at RT. Cells were washed with PBS three times and then incubated in blocking solution (PBS supplemented with 0.1% Triton-X-100 [X100, Sigma-Aldrich] and 10% normal donkey serum [017-000-121, Jackson ImmunoResearch]) for at least one hour at RT. After blocking, cells were stained with rabbit-anti-Vimentin antibody (EPR3776, Abcam) diluted 1:1000 in blocking solution overnight at 4°C. After washing three times with PBS, a donkey-anti-rabbit Alexa Flour 488 antibody (AB_2313584, Jackson ImmunoResearch) was added in a dilution of 1:500 in blocking solution. Wells were washed again with PBS and incubated in DAPI stain (1:10,000) for 10 min. Coverslips were washed again in PBS and mounted onto slides with Eukitt quick-hardening mounting medium (03989, Sigma-aldrich). Slides were dried overnight before picture caption using an Olympus CKX53 and cellSens software (Olympus). Percentage of vimentin-positive cells was determined using Fiji cell counter plugin (Schindelin et al. 2012).

### Primary Sertoli cell culture

24-well plates were coated with a DSA-lectin (L2766, Sigma Aldrich) solution (5 µg/ml in 1x Hank’s balanced salt solution [HBSS, Gibco]) for at least 1 hour at 37°C. Plates were washed twice with 1xHBSS before use. PND14 testes were enzymatically digested and resuspended in medium (DMEM high glucose [Sigma] supplemented with 0.1% bovine serum albumin [BSA, Sigma], 1x GlutaMAX [Gibco], 1x Minimum Essential Medium Non-essential Amino Acids [Gibco] and 1% Pen-Strep) at 800,000 cells/ml. 500 µl of cell suspension were added to DSA-lectin coated 24-wells and cells were allowed to attach for 2 hours at 32°C. Cells were incubated with a hypotonic solution (0.3xHBSS) for 1-2 minutes at RT to remove germ cells, washed with 1xHBSS to eliminate debris and new medium was added. Cells were left undisturbed for 24 hours before treatment.

### Serum treatment of primary Sertoli cell cultures

Cell culture medium was supplemented with 10% serum from MSUS and control adult males (batch 1), sterile-filtered using 0.22 µm PVDF filter units (Merck) and distributed to each well by individual male (1 well/mouse). After 24 hours, medium was removed and used for lactate/pyruvate assessment or snap-frozen and stored at -80°C. Cells were washed once with 1xPBS and harvested in 500 µl TRIzol (Thermo Fisher Scientific) for RNA extraction. This experiment was conducted twice in independent replicates.

### Lactate/pyruvate assessment

Conditioned medium was centrifuged at 3,200xg for 10 min at 4°C to remove debris and transferred to 10 kDa spin columns (Amicon Ultra, Merck). Proteins that may influence lactate and pyruvate level were removed from the <10 kDa flow-through containing metabolites by centrifugation at 14,000xg for 25 min at 4°C. Lactate and pyruvate were measured in the protein-depleted flow-through using assay kits (MAK064-1KT, Sigma-Aldrich) and (ab65342, Abcam) according to the manufacturer’s instructions. Each sample was run twice and fluorescence was measured on a NOVOStar Microplate reader (BMG Labtech) and averaged. For each sample, lactate/pyruvate ratio was calculated using the average lactate and pyruvate measurements of the replicates.

### ROS assessment

ROS production was measured in serum-treated primary Sertoli cultures using DCFDA/H2DCFDA-Cellular ROS Assay Kit (ab113851, Abcam) according to the manufacturer’s instructions. Fluorescence was measured immediately, then after 10, 30, and 60 minutes on a NOVOStar Microplate reader (BMG Labtech). The experiment was run in triplicates, which were averaged for each time point.

### RNA and DNA extraction

For sorted Sertoli cells obtained from adult males, RNA and DNA were extracted using the AllPrep DNA/RNA/miRNA Universal Kit (Qiagen) according to the manufacturer’s instructions. For cultured cells harvested in TRIzol (Thermo Fisher Scientific), a phenol/chloroform extraction method was used to prepare RNA.

### RNA sequencing

RNA samples were run on a Bioanalyzer (Agilent) at a concentration of 1.5 ng/µl using the eukaryote total RNA pico series II assay (Agilent) to assess RNA integrity. Libraries for RNA sequencing were prepared from 5 ng RNA/sample using the SMARTer Stranded Total RNA-Seq Kit v2 -Pico Input Mammalian (Takara) according to the manufacturer’s instructions using 12 PCR cycles for amplification. DNA concentration of libraries was determined using Qubit dsDNA HS Assay Kit, and libraries were diluted to 1.5 ng/µl, then run on a Bioanalyzer (Agilent) using the High Sensitivity DNA Assay Protocol (Agilent) for quality control. Libraries were sequenced on an Illumina NovaSeq instrument, single-end at 100 bp.

### Analyses of RNA sequencing data

Fastq files were checked for quality using FastQC (v 0.11.9) (Andrews 2010) trimmed with Trimgalore (v 0.6.5) (Krueger 2012) and pseudo-mapped with Salmon (v 1.1.0) (Patro et al. 2017) using an index file created from the GENCODE annotation of transcripts (vM23) (Frankish et al. 2019). For differential gene expression analysis, counts were normalized using the TMM method (Robinson and Oshlack 2010) and transformed with the voom method of the limma R-Package (v 3.42.2) (Ritchie et al. 2015) for linear modelling. All genes with p<0.05 were used for functional enrichment analyses using the g:GOSt function of g:Profiler (Raudvere et al. 2019), taking into account GO terms and KEGG pathways with 10-1000 annotated genes. GO terms were further simplified using Revigo (Supek et al. 2011).

### Fluidigm RT-qPCR

RNA was reverse-transcribed with miScript II RT reagents (Qiagen) using HiFlex buffer according to the manufacturer’s instructions. For high-throughput gene expression analyses, samples and primers (list of primers: S4 Table) were prepared for the Fluidigm BioMark™ HD System (Fluidigm) according to the manufacturer’s protocol. Pre-amplified cDNA samples and primers were loaded onto a 96.96 dynamic array™ (primers were loaded in duplicates) and mixed using an IFC (integrated fluidic circuits) machine (Fluidigm). Ready chips were then placed into a Fluidigm Biomark™ HD System for RT-qPCR analyses.

### Analyses of Fluidigm RT-qPCR data

Baseline correction (using linear derivative) and assessment of cycle threshold (Ct) values were performed by the BioMark HD software (Fluidigm). A list of Ct values was obtained from the BioMark output tables and ordered according to sample batch. ReadqPCR (v 1.32.0) and NormqPCR (v 1.32.0) were used for downstream data preparation, including combination of technical replicates, normalizing to the 2 most stable reference genes out of 5 (Actb, B2m, Hrpt1, Rplp0 or Vim), and deriving delta C_q_ values. Samples were normalized to the mean of control samples and log2 foldchanges were calculated.

### Determination of mitochondrial copy number variation (CNV)

DNA samples were analyzed by RT-qPCR using QuantiTect SYBR (Qiagen) on a Light Cycler II 480 (Roche): 95°C for 15 min, 45 cycles of 15 sec at 94°C, 30 sec at 55°C and 30 sec at 70°C. HK2 primers amplifying nuclear DNA were used as endogenous control and ND1 primers to amplify mitochondrial DNA (Primers list in S4 Table). Fold change of ND1 versus HK2 amplification was calculated with 2^(-delta delta CT) method and normalized to controls.

## Statistics

Student’s t-test was used to assess significance between two groups. Kenward-Roger method using R packages lmerTest (v 3.1-3) and lme4 (v 1.1-27.1) was used to assess significance for experiments run in duplicates. Outliers at a distance greater than 2.5 standard deviations from 0 were removed before analyses.

## Data availability

The RNA-sequencing datasets collected in this study are available in the Gene Expression Omnibus GSE205330.

## Acknowledgements

We thank Chiara Boscardin, Anastasia Efimova, Lola Kourouma, and Anar Alshanbayeva for assisting with the mouse work including breeding and MSUS paradigm, Andrew McDonald, Silvia Schelbert, and Alberto Corcoba for animal license and laboratory organization in Zürich, and Yvonne Zipfel and Jerome Bürki for animal care in Zürich. We are grateful for the valuable input from Pierre-Luc Germain and Deepak Kumar Tanwar for bioinformatic analyses, from Ali Jawaid for general input on study and experimental design, and Maria Dimitriu, Nancy Carullo, and Rodrigo Arzate-Mejia for proof reading. We are thankful to Niharika Obrist for technical assistance and Tao Lei and Jörg Klug from the Institute of Anatomy and Cell Biology at the University of Giessen, Germany for help with the protocol for Sertoli cell cultures. We thank the team of the cytometry facility at the University of Zurich for all FACS related issues, Aria Minder and Silvia Kobel of the Genomic Diversity Center for assistance with the fluidigm qPCR, and the team of the Functional Genomics Center Zurich for assistance with RNA sequencing. This work was funded by the University of Zurich, the ETH Zurich, the National Centre of Competence in Research (NCCR) RNA & Disease funded by the Swiss National Science Foundation (grant number 182880/Phase 2 and 205601/Phase 3), ETH grants (ETH-10 15-2 and ETH-17 13-2), and the Escher Family Fund. Illustrations in figures 1a, 3a, and 4a as well as the graphical abstract were created with BioRender.com.

## Author contributions

KMT and IMM conceived and designed the study, and wrote the manuscript. FM conducted MSUS treatment and prepared animals with the help of KMT. KMT collected and prepared Sertoli cells and serum for transcriptomic analyses and serum treatments of cell cultures. KMT prepared RNA libraries for RNA sequencing. KMT and SL prepared primary Sertoli cell cultures, and carried out serum treatment and molecular analyses. KMT analyzed the data and prepared the Figs. IMM provided conceptual support throughout the project, and raised funds to finance the project.

## Supporting information

**S1 Fig.**
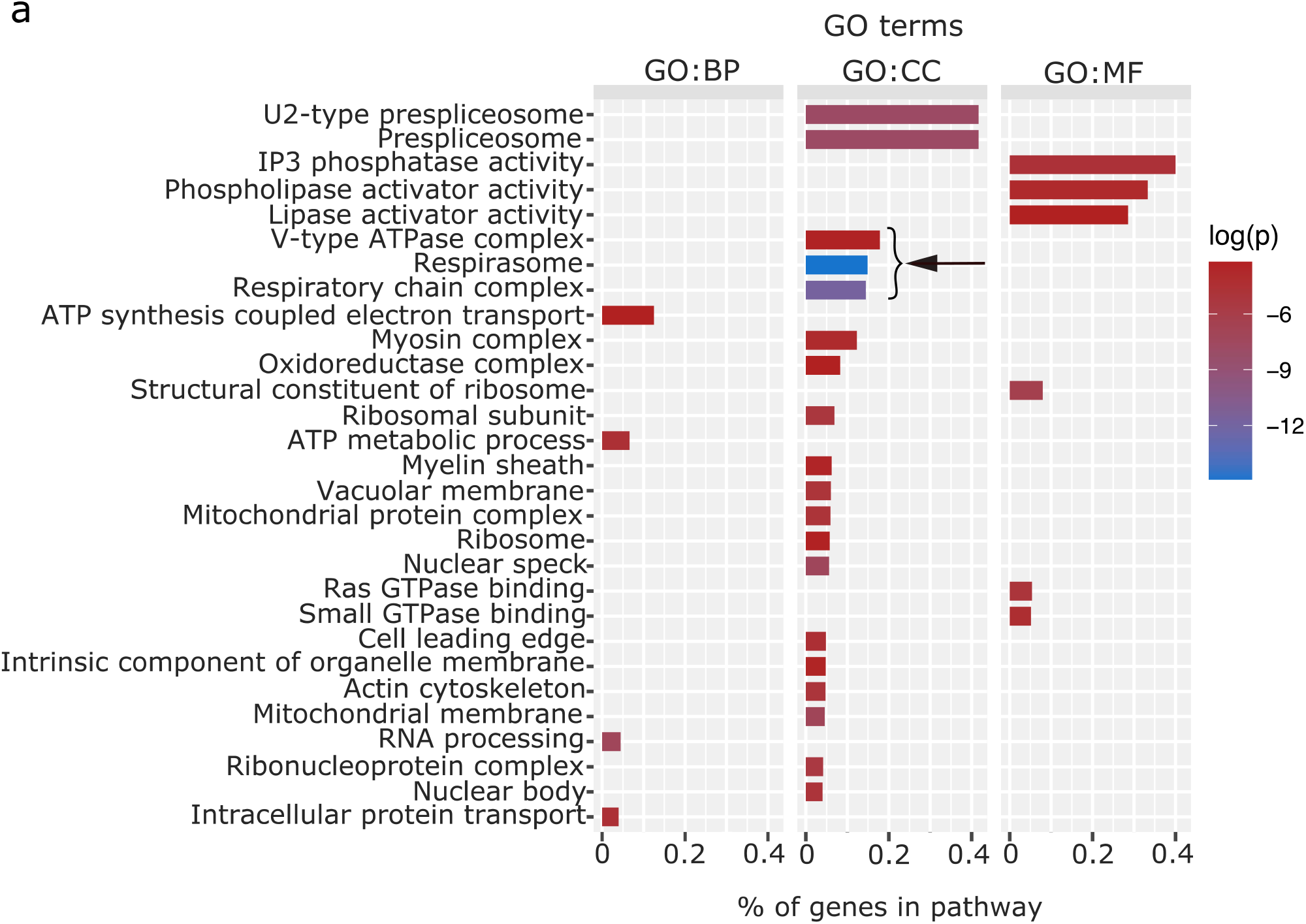
Enriched GO terms of altered genes in MSUS Sertoli cells. (a) Enriched GO pathways of significantly altered genes (p<0.05) in MSUS mouse Sertoli cells detected by RNA sequencing. Ratio of genes per pathway is given on the x-axis and log of p-value (log(p)) is indicated on a color scale. Bracket and arrow indicate mitochondrially related pathways. BP: biological process; CC: cellular component; MF: molecular function

**S2 Fig.**
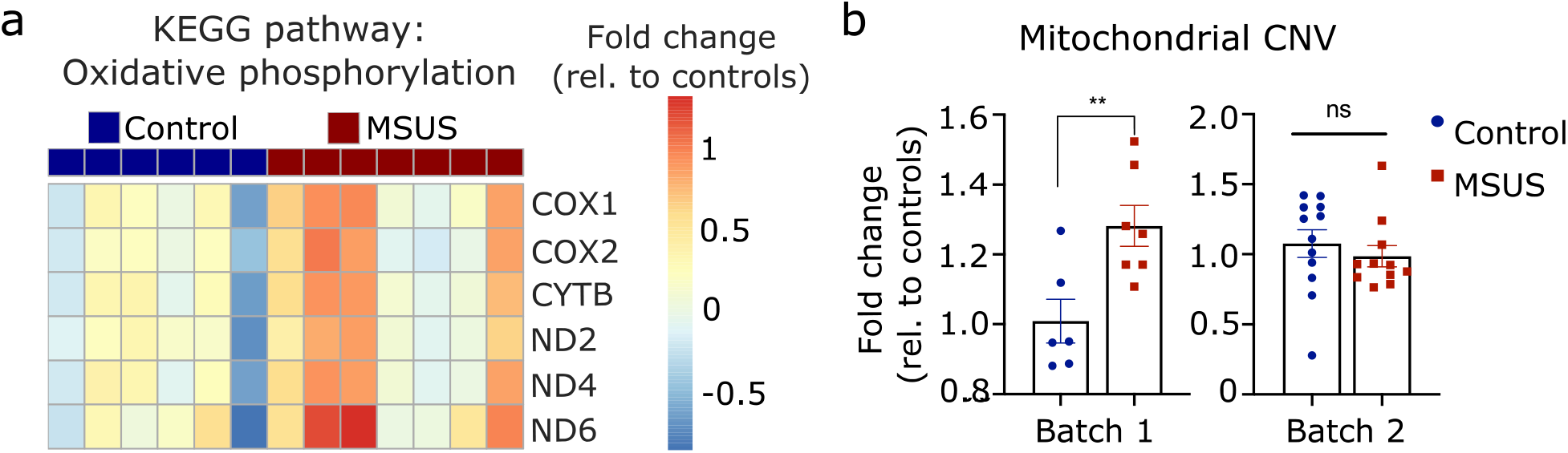
High variability in mitochondrially encoded genes and mitochondrial CNV in batch 1 Sertoli cells. (a) Heatmap of altered mitochondrially encoded genes of KEGG pathway “oxidative phosphorylation”. Fold change relative to controls is indicated in the color scale. (b) Fold changes in mitochondrial CNV of MSUS compared to control Sertoli cells of mouse batches 1 and 2. Error bars: mean ± SEM; ** = p<0.01, student’s t-test.

**S3 Fig.**
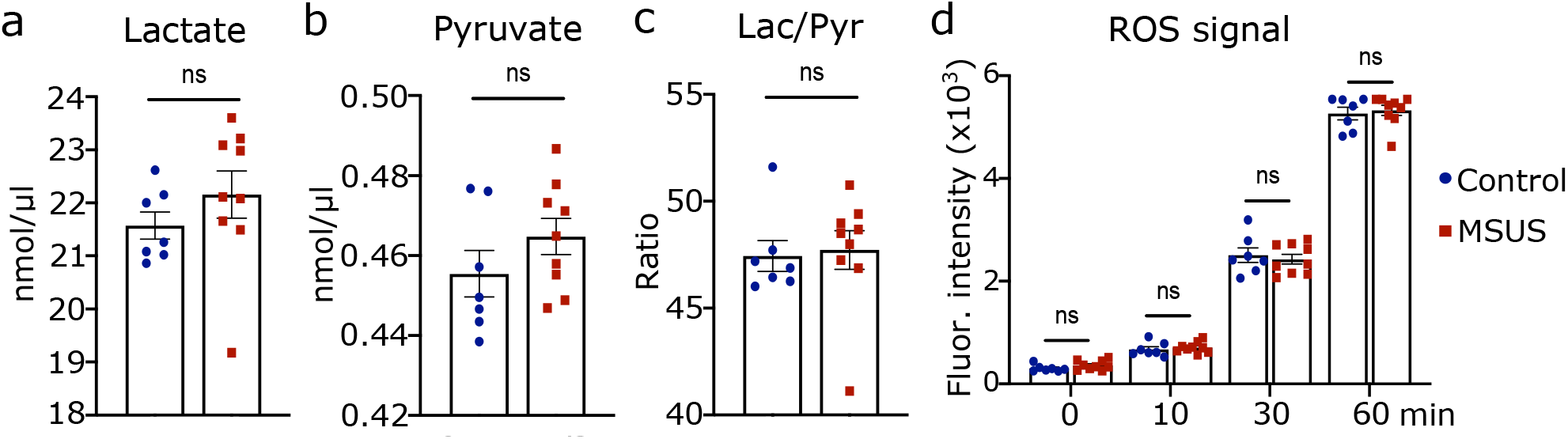
Characterization of lactate-to-pyruvate ratio and ROS production after MSUS serum exposure in primary Sertoli cells. (a) Lactate and (b) pyruvate levels (in nmol/µl) in primary Sertoli cell medium after 24h of serum exposure. (c) Ratio of lactate to pyruvate and (d) ROS fluorescent signal in primary Sertoli cells after serum exposure. Controls, n=7; MSUS, n=9 for all graphs; error bars: mean ± SEM; ns=not significant, student’s t-test.

**S1 Table.** Counts of vimentin-positive cells in isolated FACS populations. (A) Raw counts of each isolated fraction (before FACS, haploid, diploid, tetraploid, high FSC, low FSC, high APC and low APC) for 4 individual mice (4 replicates). Always at least 3 individual pictures per replicate were taken and counted. (B) Total number of cells counted for each replicate and each isolated fraction. (C) Percentages of vimentin positive cells out of all DAPI positive cells for each replicate and isolated fraction. Weighted average and weighted standard deviation (SD) were calculated from percentages.

| A | RAW COUNTS | Before |  | Haploid |  | Diploid |  | Tetraploid |  | highFSC |  | lowFSC |  | highAPC |  | lowAPC |  |
| --- | --- | --- | --- | --- | --- | --- | --- | --- | --- | --- | --- | --- | --- | --- | --- | --- | --- |
|  |  | VIM | DAPI | VIM | DAPI | VIM | DAPI | VIM | DAPI | VIM | DAPI | VIM | DAPI | VIM | DAPI | VIM | DAPI |
| Replicate 1.1 |  | 1 | 28 | 0 | 34 | 7 | 32 | 1 | 17 | 13 | 31 | 0 | 25 | 10 | 11 | 1 | 9 |
| Replicate 1.2 |  | 0 | 18 | 0 | 25 | 3 | 15 | 0 | 50 | 5 | 21 | 1 | 15 | 5 | 6 | 0 | 10 |
| Replicate 1.3 |  | 0 | 23 | 0 | 12 | 1 | 20 | 2 | 60 | 6 | 19 | 3 | 40 | 5 | 7 | 0 | 10 |
| Replicate 1.4 |  |  |  |  |  |  |  |  |  |  |  |  |  | 5 | 6 | 0 | 17 |
| Replicate 1.5 |  |  |  |  |  |  |  |  |  |  |  |  |  | 10 | 12 | 0 | 13 |
| Replicate 1.6 |  |  |  |  |  |  |  |  |  |  |  |  |  | 9 | 10 |  |  |
| Replicate 2.1 |  | 0 | 20 | 0 | 16 | 1 | 10 | 1 | 16 | 27 | 31 | 0 | 29 | 53 | 56 | 1 | 33 |
| Replicate 2.2 |  | 1 | 21 | 0 | 23 | 3 | 31 | 0 | 22 | 13 | 16 | 0 | 23 | 44 | 46 | 1 | 25 |
| Replicate 2.3 |  | 0 | 23 | 0 | 13 | 3 | 11 | 0 | 33 | 13 | 16 | 2 | 12 | 26 | 28 | 0 | 19 |
| Replicate 2.4 |  |  |  |  |  | 2 | 10 |  |  |  |  | 1 | 23 |  |  |  |  |
| Replicate 3.1 |  | 0 | 38 | 0 | 19 | 3 | 19 | 0 | 41 | 4 | 28 | 0 | 32 | 9 | 10 | 2 | 29 |
| Replicate 3.2 |  | 1 | 36 | 0 | 13 | 3 | 19 | 2 | 50 | 5 | 15 | 1 | 26 | 6 | 8 | 0 | 16 |
| Replicate 3.3 |  | 1 | 48 | 0 | 13 | 2 | 14 | 1 | 61 | 2 | 9 | 0 | 29 | 10 | 11 |  |  |
| Replicate 3.4 |  |  |  | 0 | 48 | 9 | 41 |  |  | 3 | 17 | 1 | 29 | 24 | 30 | 1 | 22 |
| Replicate 3.5 |  |  |  |  |  |  |  |  |  | 5 | 20 |  |  | 12 | 13 |  |  |
| Replicate 4.1 |  | 1 | 31 | 0 | 12 | 2 | 20 | 0 | 63 | 3 | 17 | 1 | 36 | 15 | 17 | 0 | 24 |
| Replicate 4.2 |  | 1 | 21 | 0 | 13 | 4 | 26 | 1 | 64 | 2 | 15 | 3 | 41 | 16 | 18 |  |  |
| Replicate 4.3 |  | 1 | 28 | 0 | 15 | 2 | 31 | 0 | 36 | 1 | 10 | 3 | 21 | 16 | 18 | 1 | 15 |
| Replicate 4.4 |  | 0 | 29 | 0 | 14 | 2 | 19 |  |  | 1 | 8 |  |  | 16 | 17 | 0 | 23 |

| B | #CELLS COUNTED | Before | Haploid | Diploid | Tetrapl. | highFSC | lowFSC | highAPC | lowAPC |
| --- | --- | --- | --- | --- | --- | --- | --- | --- | --- |
| Replicate 1 |  | 69 | 71 | 67 | 127 | 71 | 80 | 52 | 59 |
| Replicate 2 |  | 64 | 52 | 62 | 71 | 63 | 87 | 130 | 77 |
| Replicate 3 |  | 122 | 93 | 93 | 152 | 89 | 116 | 72 | 67 |
| Replicate 4 |  | 109 | 54 | 96 | 163 | 50 | 98 | 70 | 62 |
| TOTAL |  | 364 | 270 | 318 | 513 | 273 | 381 | 324 | 265 |

**S1 Table. Counts of vimentin-positive cells in isolated FACS populations.**
| C | PERCENTAGES | Before | Haploid | Diploid | Tetrapl. | highFSC | lowFSC | highAPC | lowAPC |
| --- | --- | --- | --- | --- | --- | --- | --- | --- | --- |
| Replicate 1 |  | 0.014 | 0.000 | 0.164 | 0.024 | 0.338 | 0.050 | 0.846 | 0.017 |
| Replicate 2 |  | 0.016 | 0.000 | 0.145 | 0.014 | 0.841 | 0.034 | 0.946 | 0.026 |
| Replicate 3 |  | 0.016 | 0.000 | 0.183 | 0.020 | 0.213 | 0.017 | 0.847 | 0.045 |
| Replicate 4 |  | 0.028 | 0.000 | 0.104 | 0.006 | 0.140 | 0.071 | 0.900 | 0.016 |
| weighted Average |  | 0.0192 | 0.0000 | 0.1478 | 0.0156 | 0.3773 | 0.0420 | 0.8981 | 0.0264 |
| weighted SD |  | 0.0063 | 0.0000 | 0.0363 | 0.0081 | 0.3036 | 0.0241 | 0.0508 | 0.0132 |

**S2 Table.**
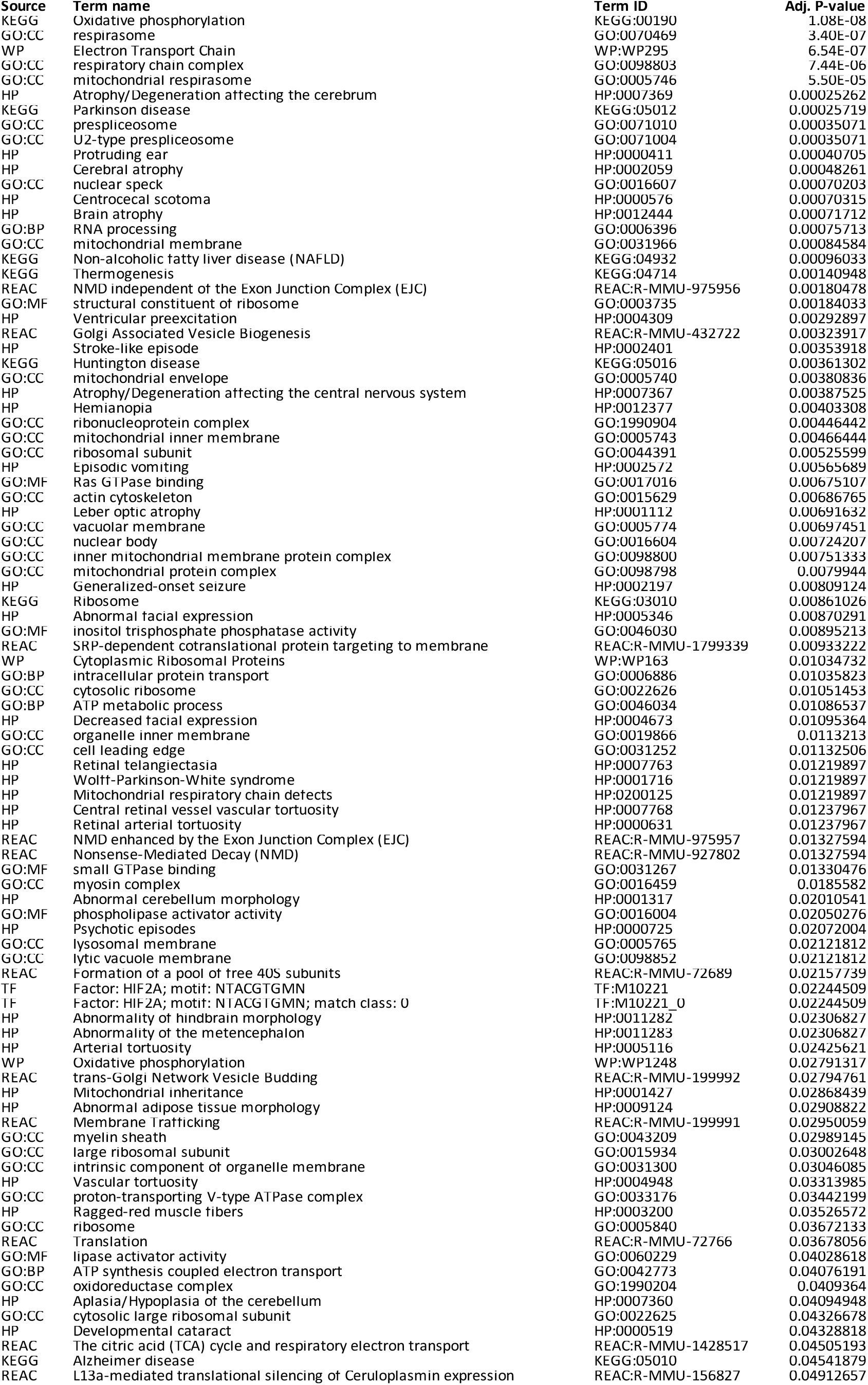
Summary of the over-representation analyses of significantly altered genes. All significantly altered genes (p<0.05) in response to MSUS were used for over-representation analyses using Gprofiler. The table shows the source, term name, term ID and adjusted p-value for each significant pathway.

**S3 Table.** Information on mice used for experiments. (A) Mice used for RNA sequencing of Sertoli cells (Batch 1). (B) Mice used for Fluidigm RT-qPCR of Sertoli cells (Batch 2). (C) Mice used for serum collection for in vitro experiments. The tables contain information on the number (ID), group, cage, and litter cage.

| A | Number | Group | Cage | Littercage |
| --- | --- | --- | --- | --- |
|  | 1 | MSUS | 11 | 576 |
|  | 5 | MSUS | 15 | 606 |
|  | 6 | MSUS | 15 | 589 |
|  | 9 | CTRL | 3 | 577 |
|  | 10 | CTRL | 3 | 583 |
|  | 11 | MSUS | 17 | 589 |
|  | 12 | CTRL | 3 | 590 |
|  | 13 | MSUS | 17 | 608 |
|  | 14 | CTRL | 3 | 601 |
|  | 22 | CTRL | 6 | 580 |
|  | 23 | MSUS | 20 | 600 |
|  | 24 | CTRL | 6 | 583 |
|  | 26 | MSUS | 20 | 597 |

| C | Number | Group | Cage | Littercage |
| --- | --- | --- | --- | --- |
|  | 9 | CTRL | 3 | 577 |
|  | 10 | CTRL | 3 | 583 |
|  | 11 | MSUS | 17 | 589 |
|  | 12 | CTRL | 3 | 590 |
|  | 13 | MSUS | 17 | 608 |
|  | 14 | CTRL | 3 | 601 |
|  | 15 | MSUS | 17 | 578 |
|  | 16 | CTRL | 6 | 596 |
|  | 17 | MSUS | 17 | 581 |
|  | 18 | MSUS | 13 | 576 |
|  | 19 | MSUS | 13 | 579 |
|  | 20 | MSUS | 13 | 588 |
|  | 21 | MSUS | 13 | 606 |
|  | 22 | CTRL | 6 | 580 |
|  | 23 | MSUS | 20 | 600 |
|  | 24 | CTRL | 6 | 583 |

**S3 Table. Information on mice used for experiments.** (A) Mice used for RNA sequencing
| B | Number | Group | Cage | Littercage |
| --- | --- | --- | --- | --- |
|  | 1 | Control | 50 | 6 |
|  | 2 | MSUS | 57 | 7 |
|  | 3 | Control | 50 | 18 |
|  | 9 | MSUS | 61 | 8 |
|  | 10 | Control | 42 | 1 |
|  | 11 | MSUS | 61 | 21 |
|  | 17 | Control | 51 | 6 |
|  | 18 | MSUS | 62 | 8 |
|  | 19 | Control | 51 | 27 |
|  | 25 | MSUS | 59 | 7 |
|  | 26 | Control | 45 | 1 |
|  | 27 | MSUS | 59 | 19 |
|  | 33 | Control | 48 | 2 |
|  | 34 | MSUS | 58 | 7 |
|  | 35 | Control | 48 | 18 |
|  | 41 | MSUS | 54 | 4 |
|  | 42 | Control | 43 | 1 |
|  | 43 | MSUS | 54 | 16 |
|  | 49 | Control | 47 | 2 |
|  | 50 | MSUS | 55 | 4 |
|  | 51 | Control | 47 | 18 |
|  | 57 | MSUS | 56 | 7 |
|  | 58 | Control | 44 | 1 |
|  | 59 | MSUS | 56 | 16 |

**S4 Table.**
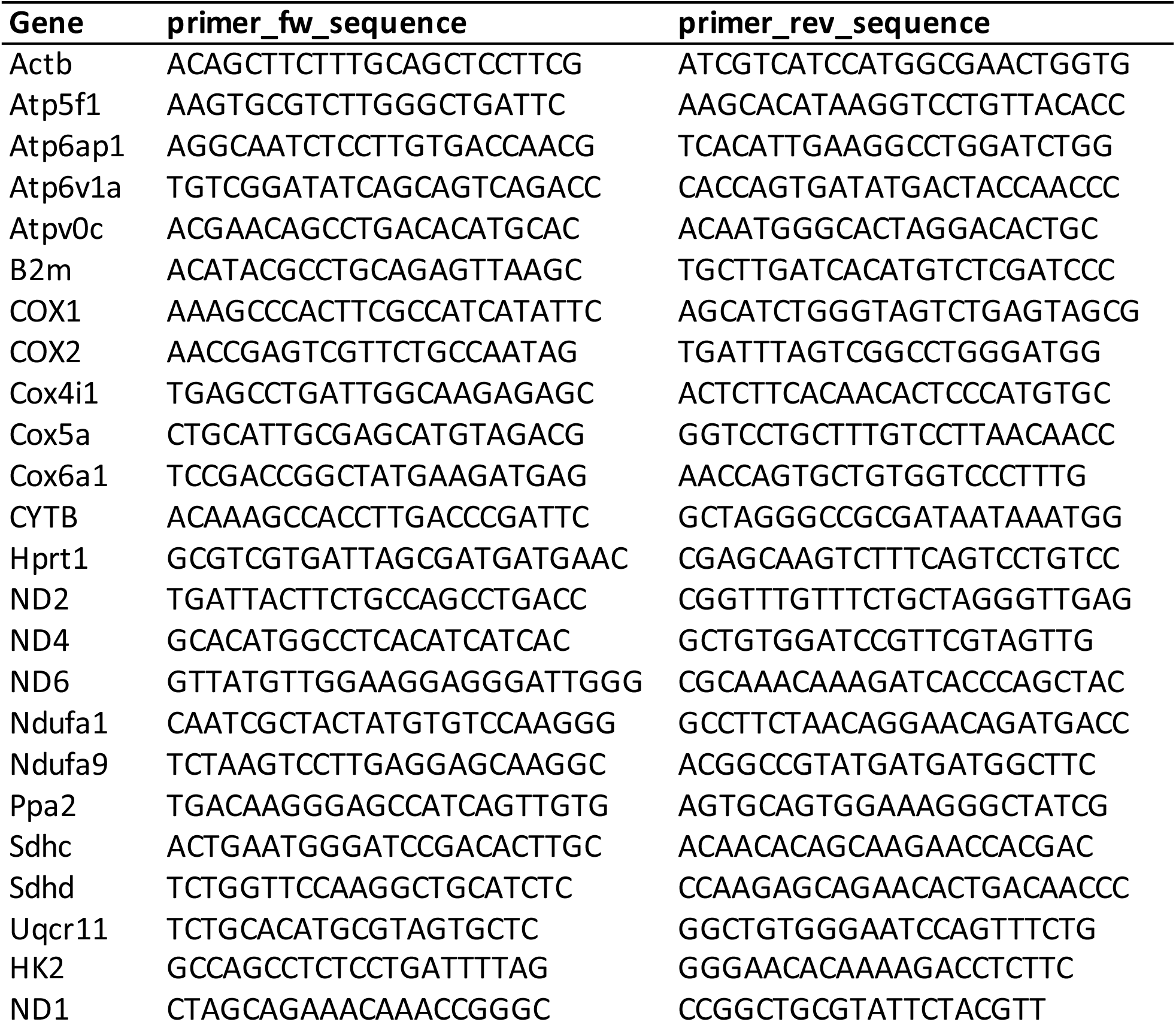
List of primer sequences. The table includes information on the target genes, forward primer and reverse primer sequences.

